# State-dependent binding of the wedge domain controls inactivation of the mechanosensitive ion channel PIEZO1

**DOI:** 10.64898/2026.01.12.699060

**Authors:** Lucas Roettger, Clement Verkest, Nadja Zeitzschel, Stefan G. Lechner

## Abstract

The mechanically activated ion channel PIEZO1 transduces membrane tension into intracellular calcium signals and is critical for a wide range of physiological processes. Recent structural and functional studies have established a detailed framework for PIEZO1 activation, but the molecular mechanisms governing its rapid inactivation remain incompletely understood. Here, we examined the contribution of the intracellular wedge domain to PIEZO1 inactivation using site-directed mutagenesis, electrophysiological recordings and MINFLUX nanoscopy. We show that wedge deletion and disruption of specific π-π and cation-π interactions between the wedge α1-helix and the pore module abolishes inactivation without impairing channel activation. Moreover, MINFLUX nanoscopy reveals that the wedge stabilizes a flat inactivated conformation of PIEZO1 and suggests that wedge dissociation is required for recovery from inactivation. Together, our data support a ball-and-chain-like mechanism with the wedge acting as a state-dependent inactivation particle that docks to the pore module to terminate channel activity during sustained mechanical stimulation.

## INTRODUCTION

PIEZO1 is a mechanically activated ion channel that converts membrane tension into rapid calcium influx, enabling cells to sense and respond to forces in their environment, such as shear stress, tissue stretch, and cell compression^1,2^. Accordingly, PIEZO1 is essential for diverse physiological processes including vascular development, blood pressure regulation, erythrocyte volume control, chondrocyte mechanotransduction, and cellular migration ^3–5^. Cryogenic electron microscopy (cryo-EM) structures revealed that PIEZO1 assembles as a homotrimer with a distinctive triskelion architecture, comprising three curved peripheral “blades” that emerge from a central pore module, which consists of an inner (IH) and an outer helix (OH) as well as a large extracellular domain known as the cap ^6–8^. The ion conduction pathway of PIEZO1 contains several gates including (from top to bottom) a cap gate, a transmembrane gate, and possibly a lateral plug gate ^9–11^. Accordingly, the choreography of gating motions and allosteric rearrangements that eventually lead to the opening of the ion-conduction pathway are complex and involve several critical steps. Cryo-EM structures obtained from different conformational PIEZO1 states^12,13^ together with in-silico modeling^14,15^, high-speed atomic force microscopy (AFM) ^16^ and MINFLUX nanoscopy of PIEZO1 in its native environment ^17–19^ have established a mechanistic framework that suggests that PIEZO1 adopts a curved conformation in the closed state, in which the blades are tilted upwards, and transitions into a flat conformation upon mechanical and chemical activation. The blades are connected to the pore module by the so-called beam – a long intracellular α-helix – which appears to act as a lever that relays the flattening motions of the blades to the pore module to promote channel activation possibly by opening the lateral plug gate ^11,20,21^. The triangular-shaped anchor domain, which resides within the plasma membrane and between the pore module and the blade, however, also appears to be involved in coupling blade flattening to the pore, as subtle changes in its geometry have profound effects on PIEZO1 sensitivity and inactivation^22^. Finally, the extracellular cap domain appears to play a central role in PIEZO1 activation. Upon flattening the cap moves downwards thereby compressing the IH– and OH-to-cap linkers, which opens the lateral cap gates such that ions can enter the central ion conduction pathway^9,13,23^.

A key property of PIEZO1 is its rapid inactivation in the presence of sustained mechanical stimulation, which appears to be essential for normal physiological function as mutations that slow down inactivation have been linked to human diseases such as hereditary stomatocytosis and xerocytosis ^3,4,24–28^. Moreover, modulation of inactivation appears to be an effective strategy to fine-tune and adjust PIEZO1-function in a cell type-specific manner by increasing cumulative calcium entry and thereby potentiating downstream signaling cascades. Thus, numerous PIEZO1 protein interaction partners, including TMEM150c ^29^, CADM1^30^, MDFIC and MDFI^31^ as well as lipids, such as ceramide ^32^, poly-unsaturated fatty acids^33^ and possibly phosphatidylinositol 4,5-bisphosphate ^34^ were shown to slow inactivation. The choreography of the allosteric rearrangements that cause inactivation and hence the mechanism by which it is altered by the afore-mentioned interactions are, however, only poorly understood. Seminal work by the Grandl lab ^35–37^ together with a recent study from Liu and colleagues ^9^ suggest that the cap domain plays a central role in inactivation. Thus, replacing specific subdomains in the cap of PIEZO1 with the equivalent domains of PIEZO2 confers rapid PIEZO2-like inactivation to PIEZO1^35,36^. Moreover, Liu et al. proposed that the flexible linkers that connect the cap to the inner and outer helix of the pore domain may serve as compressive springs that store energy upon channel flattening and subsequently release it to inactivate the channel by closing the cap gate^9^. There is, however, compelling evidence suggesting that inactivation is more complex and involves additional allosteric mechanisms. Lewis et al., for example, described two kinetically distinct inactivated states^37^, Zheng et al. found a hydrophobic inactivation gate in the inner pore ^10^ and Wu and colleagues discovered a positively charged residue in the inner helix that controls voltage-sensitivity of inactivation^36^. Moreover, disease-causing mutations that slow inactivation are not restricted to the cap and the IH but were also found throughout various other domains of PIEZO1 including the blades, the beam and the anchor^26^.

Here we examined the role of the wedge domain in PIEZO1 inactivation. The wedge domain is a short intracellular domain (32 amino acids) comprising two α-helices (α1 and α2) that are linked to the C-terminus of the beam and the N-terminus of the clasp via long flexible linkers (73 and 22 amino acids; Fig. 1A, fig. S1). An intriguing observation regarding the wedge, which sparked our interest in this domain, was that the structure of the wedge had not been resolved by any published cryo-EM structure of PIEZO1 in the curved conformation ^6–8^ and the intermediate flat conformation, potentially representing an open state ^12^ (Fig. 1A and fig. S2A), suggesting high mobility of the wedge in the closed state. In the flat inactive state, however, the wedge was resolved and appears to dock between the clasp, outer helix, and anchor domains ^13^ – a region enriched for binding sites for regulatory proteins and lipids. Thus, interaction partners such as E-cadherin^38^, SERCA^39^, TMEM150C^29^, MDFIC^31^, and phosphoinositides^34^ bind within or adjacent to this region, highlighting it as a structural hub for tuning PIEZO1 function. In this study, we investigated the role of the wedge domain in PIEZO1 function using site-directed mutagenesis and electrophysiology, complemented by MINFLUX super-resolution microscopy.

**Figure 1.**
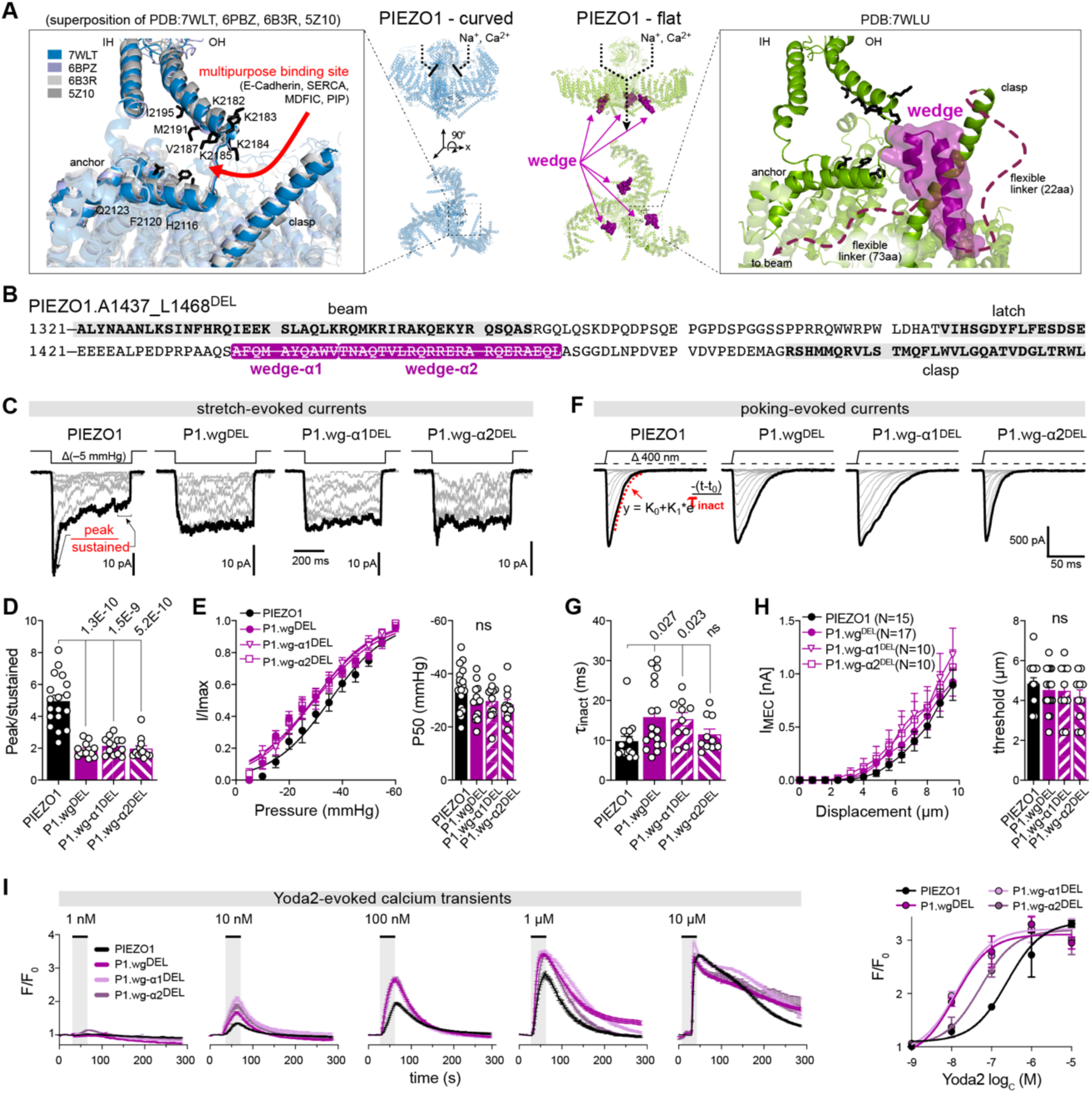
| Wedge deletion abolishes inactivation of PIEZO1. **(A)** Superposition of the indicated curved (left) and flat (right) cryo-EM structures of PIEZO1. Note, the wedge (purple) is absent from all curved structures but was reolved in close proximity to a putative multipurpose binding site in the cavity between the anchor and the outer helix (OH). **(B)** Partial amino acid sequence of PIEZO1 highlighting key domains (beam, latch, wedge and clasp). **(C)** Representative examples of stretch-evoked currents recorded from N2a-P1KO cells expressing PIEZO1 or full and partial wedge deletion mutants. **(D)** Comparision of peak/sustained ratios of the indicated stretch-evoked currents using Kruskal-Wallis test. **(E)** Normalized pressure-response curves of the indicated channels (left) and comparison of P50 values (right) using one-way ANOVA with Dunnett’s multiple comparisons test. **(F)** Representative examples of poking-evoked currents recorded from N2a-P1KO cells expressing PIEZO1 or full and partial wedge deletion mutants. **(G)** Comparison of inactivation time constants of poking-evoked currents, determined by exponential decay fits (see F), using Kruskal-Wallis-test with Dunn’s multiple comparison test. **(H)** Displacment–response curves (left) and comparison of mechanical activation thresholds of poking-evoked currents mediated by the indicated channels **(I)** Average time-courses of Yoda2-evoked calcium transients (left) measured as F/F0 (= GCamp8 fluorescence intensity at time t divided by the average GCamp8 intensity during the first ten frames) and dose-response curves of Yoda2 for the indicated channel mutants. All bars in D-H represent means ± s.e.m., with white circles showing individual values of biological replicates and P-values being provided above bars.

## RESULTS

### The wedge domain controls inactivation of PIEZO1

To examine the role of the wedge domain in PIEZO1 function, we generated mutant channels that lack the full wedge (PIEZO1.A1437_L1468^DEL^), hereafter referred to as P1.wg^DEL^, as well as partial wedge deletions that lack either the α1– or the α2-helix of the wedge (P1.wg-α1^DEL^ and P1.wg-α2^DEL^; Fig. 1B) and characterized mechanically-evoked currents as well as chemically-induced calcium transients mediated by these channels using patch-clamp electrophysiology and GCamp8 calcium imaging, respectively. The mutant channels were expressed in Neuro2a-PIEZO1 knockout cells (N2a-P1KO)^40^ and mechanotransduction currents were evoked by pressure-induced membrane stretch in the cell-attached patch-clamp configuration (i.e. pressure-clamp technique) or by focal mechanical stimulation with a small glass rod in the whole-cell patch-clamp mode (poking technique^41^, see reference ^42^ for comparison of the two techniques). All three mutant channels were trafficked to the plasma membrane (Fig. S2B) and produced robust mechanically activated currents in the pressure-clamp and poking assay. Notably, while the three mutants were indistinguishable from full-length PIEZO1 with regards to current amplitudes, activation thresholds and single channel conductance, they exhibited significantly slower inactivation during sustained mechanical stimulation (Fig. 1C-H, fig. S2C-E). Thus, the ratios of peak current amplitudes to sustained current amplitudes in pressure-clamp recordings (see Fig. 1C-D) significantly decreased from 4.95 ± 0.37 (PIEZO1; mean ± s.e.m; N = 18) to 1.89 ± 0.12 (P1.wg^DEL^; N = 13, P = 3.67E-6; Kruskal-Wallis test), 2.15 ± 0.15 (P1.wg-α1^DEL^; N = 13, P = 2.15E-4) and 1.97 ± 0.19 (P1.wg-α2^DEL^; N = 13, P = 9.63E-6) upon full or partial deletion of the wedge domain. Likewise, the inactivation time constants of poking-evoked currents, determined by exponential decay fits (Fig. 1F–G), increased from 9.8 ± 1.2 ms (PIEZO1, N = 15) to 15.8 ± 2.1 ms (P1.wg^DEL^; N = 17, P = 0.027; Kruskal-Wallis test) and 15.3 ± 1.6 (P1.wg-α1^DEL^; N = 10, P = 0.024; Kruskal-Wallis test) in the absence of the full wedge and the wedge-α1 helix but not upon deletion of the wedge-α2 helix (11.5 ± 1.4 ms; N = 10, P = 0.83; Kruskal-Wallis test). Consistent with the slowed inactivation, which prolongs the time the channels spend in the open state and thus allows more calcium to enter the cell, calcium transients triggered by the selective PIEZO1 activator Yoda-2 were also larger and detectable at lower Yoda-2 concentrations in N2a cells transfected the wedge-mutants as compared to full-length PIEZO1. Similar to poking-evoked currents, deletion of the wedge-α1 helix had a more pronounced effect on calcium transients than wedge-α2 deletion.

Together, these data suggest that the wedge domain plays a crucial role in controlling the inactivation of PIEZO1 and suggest that this modulatory effect is predominantly mediated by the wedge α1-helix.

### The wedge binds to the anchor-OH-linker via hydrophobic interactions

To test this hypothesis, we next analyzed possible non-covalent interactions between the wedge and adjacent domains in the flattened PIEZO1 structure (PDB:7WLU)^13^ using the RING4.0 platform^43^. This analysis revealed that tyrosine 1442 (Y1442) and tryptophane 1445 (W1445) of the wedge α1 helix are in close proximity to Y2175 and K2184 of the anchor-OH-linker and thus possibly dock the wedge to the pore module via cation-π and π–π interactions, respectively (Fig. 2A). Moreover, this analysis suggested that R1456 of the wedge α2-helix forms a hydrogen bond with Q1510 of the nearby clasp domain to further stabilize wedge binding. To examine the possible role of these non-covalent interactions in PIEZO1 inactivation, we generated PIEZO1 single and double mutants in which the afore-mentioned amino acids were substituted by alanines. Strikingly, all mutants that altered possible interactions between the wedge α1-helix and the anchor-OH-linker, significantly (except K2184A) slowed down inactivation (peak/sustained ratio, PIEZO1 = 4.95 ± 0.37, N = 18; P1.Y1442A = 1,79 ± 0.14, N = 12; P1.W1445A = 2.09 ± 0.17, N =1 3; P1.Y1442A.W1445A = 1.91 ± 0.07, N = 11; P1.Y2175A = 1.84 ± 0.08, N = 12; P1.K2184A = 2.86 ± 0.15, N = 8; P1.Y2171A.K2184A = 1.59 ± 0.03, N = 11; P1.R1456A = 8.29 ± 0.14, N = 8), whereas disrupting the possible wedge-clasp interaction by mutating R1456 did not have any effect. Interestingly, unlike full and partial deletion of the wedge, which did not alter PIEZO1 sensitivity, abrogation of possible wedge–anchor-OH-linker interactions also shifted the pressure-response curves towards less negative pressures (Fig. 2E) and thus significantly reduced the mean pressures required for the half-maximal activation of the channels (P_50_) from 34.6 ± 1.9 mmHg in PIEZO1 (N = 18) to values around ∼25 mmHg (P1.Y1442A = 24.2 ± 1.6 mmHg, N = 12; P1.W1445A = 25.4 ± 1.2 mmHg, N =13; P1.Y1442A.W1445A = 26.7 ± 1.8 mmHg, N = 11; P1.Y2175A = 27.6 ± 0.7 mmHg, N = 12; P1.K2184A = 27.5 ± 2.3 mmHg, N = 8; P1.Y2171A.K2184A = 20.5 ± 1.3 mmHg, N = 11; P1.R1456A = 34.5 ± 2.4, N = 8; Fig. 2E).

**Figure 2.**
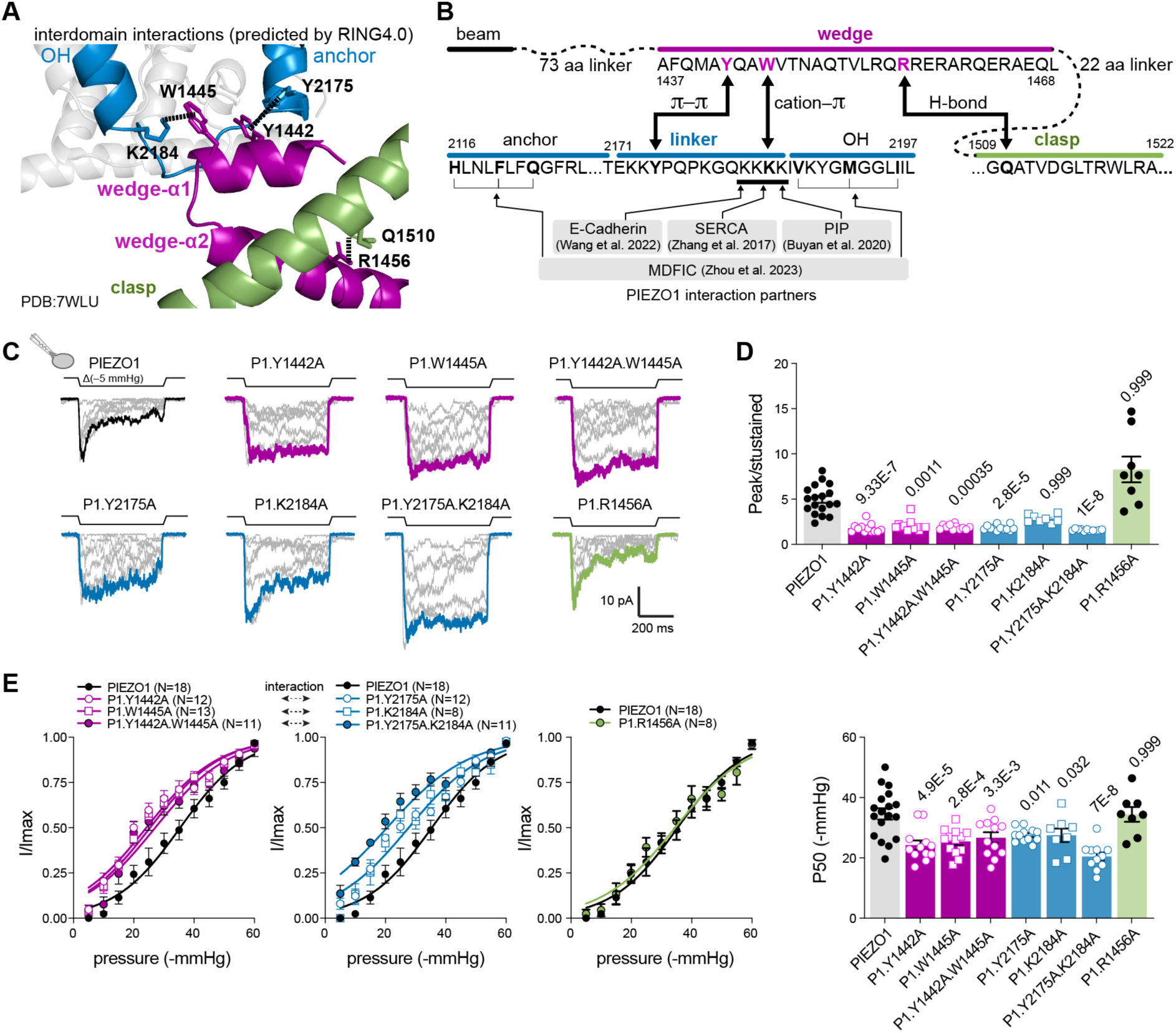
| Non-covalent interactions between the wedge α1 helix and the anchor-OH linker are required for wedge docking. (**A**) Close-up of the anchor-OH linker region in the flat PIEZO1 structure (PDB: 7WLU) with putative interactions with the wedge as predicted by RING4.0. **(B)** Partial amino acid sequence of PIEZO1 showing the wedge, anchor, linker, OH and clasp domains and amino acids engaged in the putative interdomain interactions and interactions with other proteins highlighted in bold letters. **(C)** Representative examples of stretch-evoked currents recorded from N2a-P1KO cells expressing PIEZO1 or the indicated point mutants. **(D)** Comparision of peak/sustained ratios of the indicated stretch-evoked currents using Kruskal-Wallis with Dunn’s multiple comparison test. **(E)** Normalized pressure-response curves of the indicated channels fitted with Boltzman sigmoidal functions (left) and comparison of P50 values (pressure required for half-maximal channel activation, right) using one-way ANOVA with Dunnett’s multiple comparisons test. All bars in D and F represent means ± s.e.m., with white circles showing individual values of biological replicates and P-values being provided above bars.

In summary, our patch-clamp recordings from full and partial wedge deletions as well as PIEZO1 point mutants together with previous cryo-EM data suggest that – in the flat (open) state – the wedge binds to the anchor-OH-linker via cation-π and π–π interactions, thereby promoting rapid inactivation of PIEZO1.

### PIEZO1 instantly recovers from inactivation in the absence of the wedge

To test if the lack of current decay during sustained stimulation of the P1.wg^DEL^ mutant, indeed resulted from a lack of inactivation (i.e. transition from the open to the inactivated state) rather than altered deactivation (open to closed) or adaptation (e.g. patch deterioration, seal creep ^44^, cytoskeletal rearrangements ^45^), we next applied two identical and supramaximal test pulses (−60 mmHg) at an interval of 5 seconds with the second test pulse being preceded by a 500 ms conditioning pre-pulse to –20 mmHg, which partially activates PIEZO1 and allows currents to decay back to baseline levels (Fig. 3A). In cells expressing full-length PIEZO1 the second test pulse (+ pre-pulse) consistently elicited smaller currents (I_+prepulse_/I_–prepulse_ = 0.45 ± 0.05, N = 11) demonstrating that a considerable fraction of PIEZO1 channels indeed transition into an inactivated state during sustained mechanical stimulation in which they are reluctant to open in response to subsequent mechanical stimuli. By contrast, P1-wg^DEL^ currents reached almost the same peak amplitudes after application of the conditioning pre-pulse currents (I_+prepulse_/I_–prepulse_ = 0.85 ± 0.05, N = 16), suggesting that channels do not inactivate in the absence of the wedge and immediately return to a closed and “willing” state upon stimulus removal.

**Figure 3.**
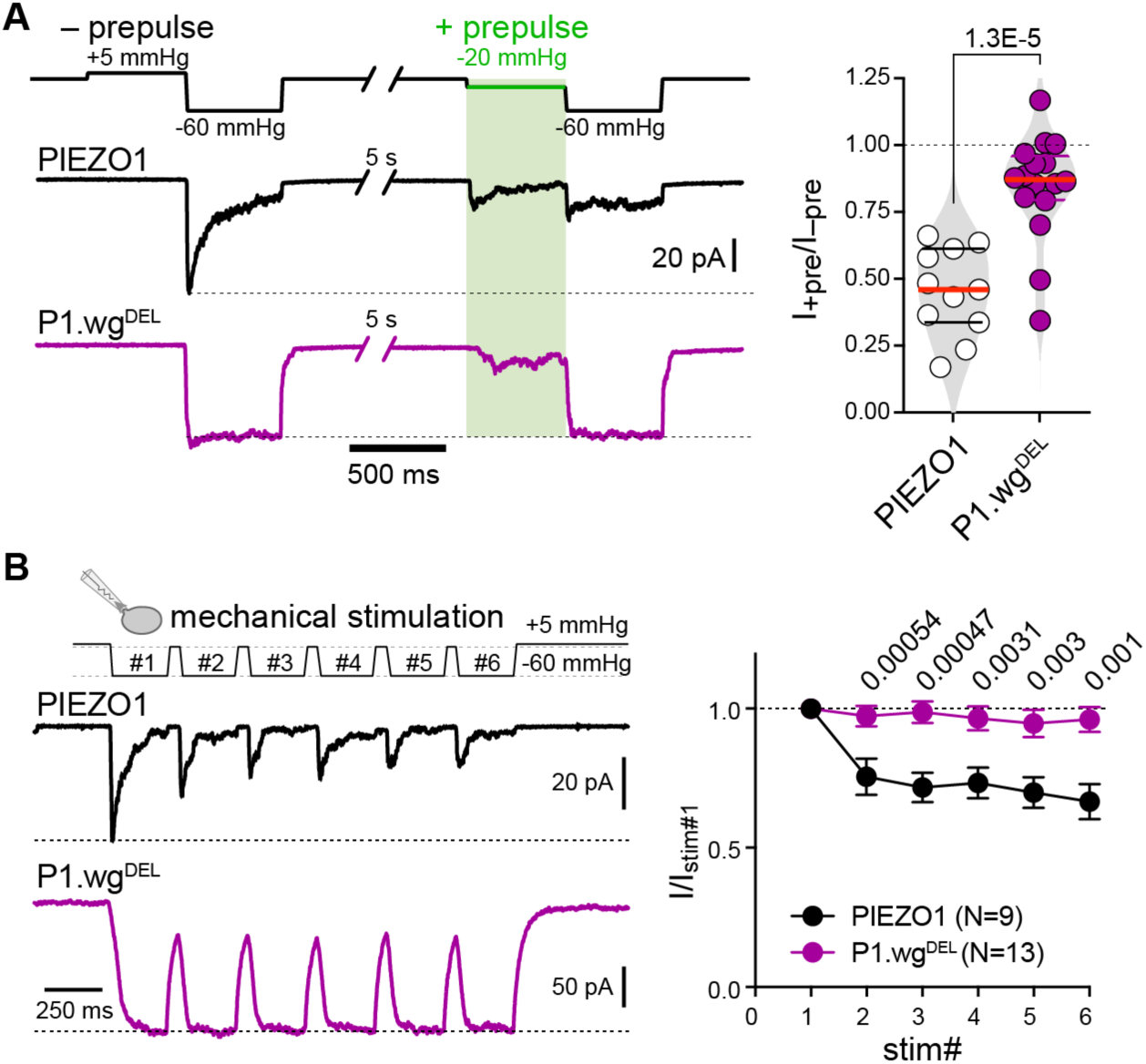
| PIEZO1 instantly recovers from inactivation in the absence of the wedge. (**A**) Representative example traces of PIEZO1 (top) and P1-wg^DEL^ (bottom) currents evoked with (right) or without (left) a conditioning pre-pulse to –20 mmHg and comparison of the ratios of peak current amplitudes elicited with and without conditioning pre-pulse. Circles show ratios of biological replicates, medians are indicated by horizontal red lines and were compared using two-sided Student’s upaired t-test (P = 1.36E-5). **(B)** Example traces of PIEZO1 (top left) and P1.wg^DEL^-mediated (bottom, left) currents elicited by repetitive mechanical stimulation. Magnitude of responses normalized to the first response of PIEZO1 (black) and P1.wg^DEL^ (purple). Symbols represent means ± s.e.m. and p-values of multiple unpaired comparison t-tests are provided above.

We next considered recovery from inactivation. Previous studies have shown that PIEZO1 remains in an inactivated state after removal of the mechanical stimulus and that full recovery from inactivation can take up to ten seconds^37^. To test if wedge deletion alters recovery from inactivation, we applied six consecutive and supramaximal pressure pulses (magnitude = –60 mmHg; duration = 250 ms) with varying time intervals (50 ms, Fig. 3A, 100 ms, 200 ms, 500 ms and 1000 ms; Fig. S2A) in cell-attached recordings and measured the decay of peak current amplitudes. Consistent with previous reports^37^, full-length PIEZO1 peak current amplitudes got smaller with each stimulus – even at the longest test interval of 1000 ms – demonstrating that the channels only slowly recover from inactivation (Fig. 3B and fig. S3A). By contrast, the peak amplitudes of P1.wg^DEL^ currents did not decay at all and were identical throughout the entire stimulation protocol even when the consecutive test pulses were applied at extremely short intervals of 50 ms (Fig. 3B and fig. S3A).

### PIEZO1 remains in a flat conformation after inactivation

It is well-established that PIEZO1 adopts a curved conformation in the closed state and transitions into a flat conformation upon mechanical and chemical activation^14,16,19,46^. The allosteric mechanisms and conformational changes underlying inactivation are, however, only poorly understood. Previous studies from others and us that examined the distribution of conformational states of PIEZO1 in its native environment – using MINFLUX nanoscopy ^18,19,46^ and cryo-light microscopy ^47^ – demonstrated that a large fraction of PIEZO1 channels (∼30-40%) are captured in a flat conformation even in the absence of mechanical stimulation. Considering that PIEZO1 only shows little spontaneous activity in the absence of mechanical stimulation, we hypothesized that these flat channels represent a flat inactivated conformation that might be stabilized by docking of the wedge.

To test this hypothesis, we utilized 3D-MINFLUX nanoscopy in combination with DNA-PAINT, which achieves spatial resolutions in the nanometer range ^48–50^, thereby enabling us to resolve the distances between fluorophores attached to the peripheral ends of the blade domains – i.e. interblade distance (a surrogate measure for channel flattening) – and thus directly observe PIEZO1 conformation in its native environment (Fig. 4A and B). To this end, we inserted ALFA-tags^51^ into position H86 (extracellular distal end of the blade), which does not alter channel function ^18,19^, and imaged cells with 3D-MINFLUX using a DNA-PAINT docking strand-conjugated single domain anti-ALFA nanobody and an Atto655-conjugated DNA-PAINT imager strand (Fig. 4A and B and fig. S4A). This labelling system has a linkage error of only ∼2.5 nm, thereby allowing precise position estimates of the distal ends of the blades (Fig. 4A). Furthermore, the MINFLUX data obtained here had a localisation precision of less than 5 nm in all three dimension (fig. S4B). To identify triple-labelled PIEZO1 trimers in MINFLUX localisation data, we computed the Euclidian distance matrix of all localisations and searched for localisation triplets in which the distances between the individual localisations were smaller than 40 nm (i.e. the maximal physically possible distance between two ALFA-tags in the fully flattened state) and that had no other neighbouring signals within 60 nm. Since triple-labelled PIEZO1 trimers are relatively scarce, due to technical limitations of MINFLUX and limitations in labeling efficiency ^17–19^, we also analyzed the interblade distances of double labelled trimers using the same cut-off criteria. Using this approach we have previously shown that PIEZO1 conformation at rest varies between cellular compartments and is governed by cytoskeletal rigidity^18^ and have demonstrated Yoda-induced flattening of the channel^19^. In agreement with the work from Mazal and colleagues^47^ and our own previous work ^18,19^, the interblade distance distribution of full-length PIEZO1 observed here was broad and non-uniform (Fig. 4C and D) and could be faithfully fitted with a Gaussian mixture model with six components (N=105, R^2^ = 0.937), with four broad peaks below 25.5 nm, which likely represent curved conformations with different numbers of ‘handshakes’ between the peripheral blades^18,52^, and two distinct peaks at values greater than 25.5. nm, which possibly represent the intermediate^12^ and the fully flattened^13^ conformations, repsectively (Fig. 4D). Notably, the median interblade distance measured in triple-labelled trimers was significantly reduced by 3.59 nm in the P1.wg^DEL^ mutant as compared to full-length PIEZO1 (PIEZO1 = 23.2 ± 0.6 nm, N = 105 vs. P1.wg^DEL^ = 19.6 ± 0.6 nm, N = 103, P = 8.2E-5, Mann-Whitney test; Fig. 4C). Measurements of the interblade distances of double-labelled PIEZO1 trimers showed the same difference (fig. S4C) and the interblade angles were not altered by wedge deletion, indicating that the wedge only affects blade flattening but not rotational flexibility (fig. S4D). Detailed inspection of the interblade distance distribution histogram showed that the peak representing the fully flattened conformation was almost completely absent and the peak representing the intermediate flat conformation became significantly smaller in the interblade distance distribution of P1.wg^DEL^ (purple, Fig. 4D), whereas the leftmost peak, which possibly represents the curved conformation with three handshakes (dark blue, Fig. 4D), was significantly larger. To statistically compare the two distributions, we categorized all channels with interblade distances below 25.5 nm as curved and those with interblade distances greater than 25.5. nm as flat and preformed a Fisher’s exact test. This analysis revealed that the overall proportion of channels that adpot a flat state significantly decreases from 40.9 % to 18.4 % upon deletion of the wedge (P = 0.0005, Fisher’s exact test, Fig. 4E), suggesting that the wedge normally stabilizes a flat inactivated conformation.

**Figure 4.**
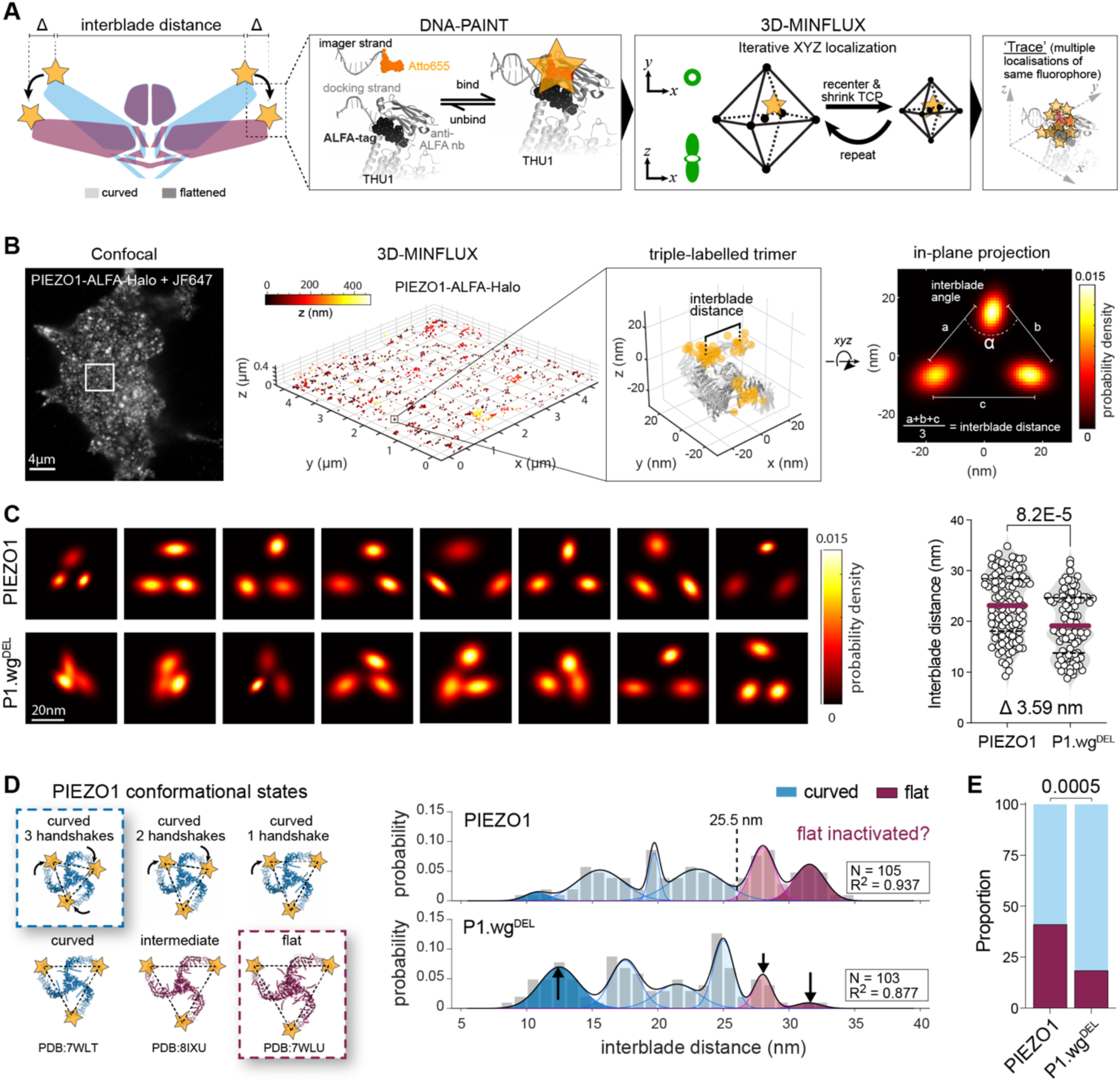
| The wedge stabilizes a flat inactivated conformation. (**A**) Cartoon depicting the overall strategy to resolve PIEZO1 conformation in its native environment by measuring interblade distances. Insets (from left to right) illustrate the labelling of PIEZO1 with ALFA tag inserted after H86 and the DNA-PAINT method (left), the 3D-MINFLUX targeted coordinate pattern used for fluorophore position determination in 3D (middle) and example of a MINFLUX ‘trace’ – i.e. series of repeated localisations of the same fluorophore molecule. **(B)** Workflow example showing confocal image of a PIEZO1-ALFA-mGL expressing N2a-P1KO cell (left) and corresponding 3D-MINFLUX localizations (second from left), colored by Z position. Inset shows a triple-labelled PIEZO1, with the traces (multiple localizations) of the three fluorophores superimposed on cryo-EM structure. The 2D in-plane projections of the 3D data were fitted with a bivariate Gaussian distribution, with their probability densities, enabling determination of the average interblade distance (right). **(C)** In-plane projections of representative trimers examples of PIEZO1 (top left) and P1.wg^DEL^ (bottom left) together with a comparison of interblade distances (right) of PIEZO1 (N=105) and P1.wg^DEL^ (N=103) using two-sided Mann-Whitney test (P = 8.2E-5) **(D)** Top views (left) of the indicated cryo-EM structures of possible closed (blue) and open (purple) states of PIEZO1. To enable depiction of interblade distances, the peripheral ends of the blade domains, which were not resolved by cryo-EM, were modelled by aligning the full-length Alphafold structure (AF-E2JF22) with cryo-EM structures. Domains originating from the Alphafold structure are shown as semi-transparent cartoons. To illustrate ‘handshaking’ between adjacent blades (top row), the peripheral blades were manually rotated counterclockwise. The right panel shows the interblade distance frequency distribution histograms of PIEZO1 (top) and P1.wgDEL (bottom) fitted with a Gaussian mixture model with six components. Peaks of putative flat comformation are shown in purple and those of curved states in blue. **(E)** Comparison of the proportions of channels adopting flat (purple) and curved (blue) conformations using Fisher’s exact test (P = 0.0005).

In summary, previous cryo-EM data together with the electrophysiological and MINFLUX data obtained here support a ball-and-chain-like inactivation mechanism in which the wedge serves as an inactivation particle that docks to the anchor-OH-linker in a state-dependent manner (i.e. only in the flat conformation) to induce inactivation and stablize a flat inactivated conformation (Fig. 5).

**Figure 5.**
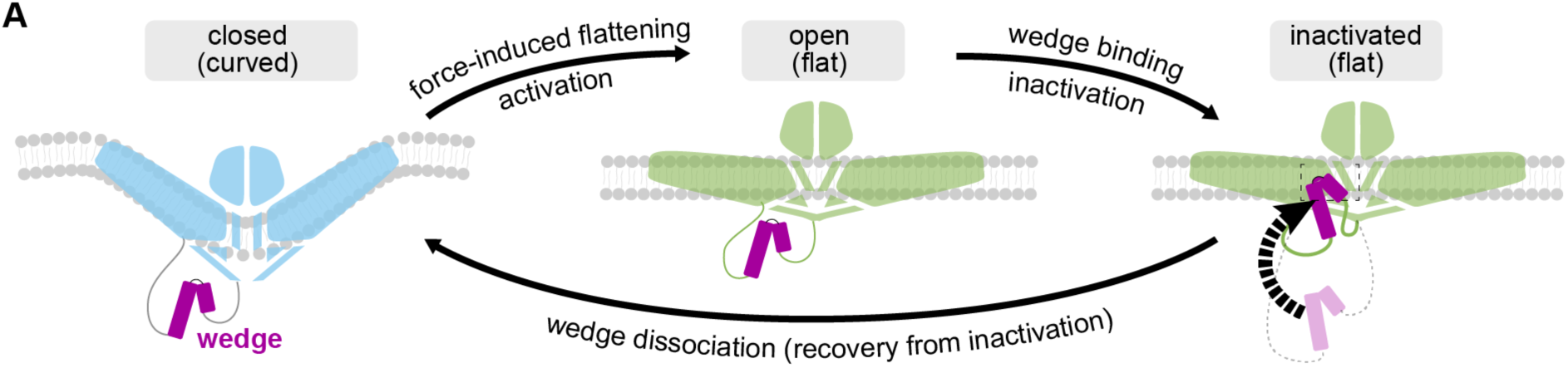
| Mechanistic model of PIEZO1 inactivation via wedge docking. (**A**) Cartoon illustrating the mechanistic model of inactivation supported by our data.

## DISCUSSION

Inactivtation is a common self-regulation mechanism of many ion channels, including voltage-gated sodium and potassium channels, some ligand-gated ion channels and not least PIEZO channels, which limits ion influx in the continuous presence of an activating stimulus and is thus essential for normal cellular function. In voltage-gated sodium and potassium channels rapid inactivation is mediated by a mechanistically conserved ball-and-chain mechanism, where a small inactivation particle that is connected to the α-subunit of the channel via a flexible polypeptide chain docks to the pore or adjacent to the pore to terminate ion influx ^53,54^. Here, we provide evidence suggesting that a similar ball-and-chain-like mechanism, where the wedge serves as the inactivation particle and the flexible linkers that connect the wedge to the clasp and the beam serve as the chain, contributes to the inactivation of PIEZO1 (Fig. 5). This inactivation model is supported by three key observations: firstly, the wedge has not been structurally resolved in any previously published cryo-EM structure of the curved (closed) PIEZO1 conformation^6–8^, whereas it was found docked to the lateral side of the pore module in the flat PIEZO1 conformation ^13^ (Fig. 1A), which strongly suggests state-dependent wedge binding. Secondly, deletion of the wedge or disruption of specific non-covalent π-π and cation-π interactions between the wedge α1-helix and the anchor–OH-linker almost completely abolished inactivation in pressure-clamp recordings without otherwise impairing PIEZO1 function (Fig. 1C, D, F and G, Fig. 2C and D). Thirdly, measurements of PIEZO1 interblade distances – i.e. a surrogate measure for channel flattening – in the presence and absence of the wedge domain, using 3D-MINFLUX nanoscopy, indicate that the wedge is required to stabilize a flat and possibly inactivated conformation (Fig. 4).

While evidence for state-dependent binding of the wedge domain is solid and comes from three different laboratories that resolved the conformation of PIEZO1 in the curved ^6–8^ and flat^13^ state using cryo-EM, the whereabouts of the wedge in the curved state and the choreography of its motions during inactivation are less clear (Fig. 1A). The secondary structure prediction algorithms PSIRPED, JNET and TRANSSEC^55–57^, together with the intrinsic disorder prediction tools IUPRED3 and PONDR(VL-XT)^58^ suggest that the wedge – irrespective of PIEZO1 conformation – adopts a stable secondary structure comprising two α-helices and that the linkers that attach the wedge to the clasp and beam are intrinsically disordered and thus probably highly flexible (fig. S1A and B). Moreover, Alphafold also assigns a defined secondary structure with two α-helices to the wedge domain in the curved state but places the wedge distant from the anchor–OH–linker where it cannot engage in non-covalent interactions with this region (AF-E2JF22) ^59^. Hence, our electrophysiological characterization of the point mutants that disrupt putative non-covalent interactions between the wedge and the anchor-OH-linker in the flat state (Fig. 2) together with the experimentally determined structures (Fig. 1A) and the in-silico predictions (fig. S1A and B) strongly suggest that the wedge together with the flexible linkers form a ball-and-chain-like inactivation system that terminates ion flux during sustained mechanical stimulation by binding to the anchor-OH-linker. We have not examined the allosteric rearrangements that eventually lead to inactivation downstream of wedge docking and can thus only speculate about this mechanism. Previous studies have demonstrated a prominent role of the cap and the inner helix in PIEZO1 inactivation ^9,10,35,36^. Hence, considering that the cap-OH spring linkers and the hydrophobic inactivation gate (L2475 and V2476), which are required for cap– and IH-dependent inactivation, respectively, are in close proximity to the wedge docking site, one possible explanation would be that wedge docking facilitates cap– and/or IH-dependent inactivation via allosteric coupling. It is, however, also possible that wedge-dependent inactivation represents a different and independent inactivation mechanism. Indeed, the wedge docking site – more generally the anchor-OH-linker – has previously been shown to be a crucial hub for the interaction with several PIEZO1 modulators that alter inactivation, including E-cadherin^38^, SERCA^39^ and PIP^34^. Moreover, recent cryo-EM structures complemented with functional patch-clamp data showed that the space between the anchor, OH and IH serves as a binding pocket for the auxiliary PIEZO1 subunit MDFIC^31^, which also controls channel inactivation, and for a yet unidentified – possibly a lipid – cofactor that facilitates full channel flattening^60^. Thus, another possible mode-of-action of the wedge is that it directly competes with the afore-mentioned modulators or indirectly blocks access to their binding sites such that they can exert stronger effects (i.e. slow inactivation) in the absence of the wedge.

We also examined the conformational changes involved in the recovery from inactivation using 3D-MINFLUX nanoscopy (Fig. 4). An intriguing observation from previous super-resolution imaging studies of PIEZO1 in its native environment by others and us^18,19,46,47^, was that an unexpectedly large fraction of PIEZO1 channels exist in the flattened conformation in the absence of mechanical stimulation, which is believed to be the main driving force for channel flattening. One possible explanation for this phenomenon is that PIEZO1 remains trapped in the flat conformation for quite some time after inactivation before returning to the curved conformation. Rare spontaneous activation events (e.g. triggered by cell-generated traction forces) could thus cause accumulation of flat – yet inactivated – channels over time, even in the absence of exogenous mechanical stimulation, which could explain the large fraction of flat channels although PIEZO1 only shows little background activity. Indeed, previous studies have shown that recovery from inactivation is very slow and may take up to ten seconds or longer ^36,37^ and traction force-induced activation of PIEZO1 has also been elegantly demonstrated using advanced live-cell imaging ^61^. Our 3D-MINFLUX data further supports this hypothesis, by showing that the interblade distribution of the full wedge deletion PIEZO1 mutant, which lacks inactivation (Fig. 1C and D) and immediately returns to a closed and “willing” state upon removal of the mechanical stimulus (Fig. 3), completely lacks the peak representing channels in the fully flattened conformation, while indicating that many more channels adopt a curved conformation (Fig. 4D and E), suggesting that wedge docking normally stabilizes the flat conformation.

Hence, we propose a refined inactivation model for PIEZO1 where the wedge, in addition to the cap and inactivation gate of the inner helix, contributes to inactivation by binding to the anchor-OH-linker in a ball-and-chain-like manner. We further propose that wedge docking stabilizes the flat inactivated conformation, and that dissociation of the wedge is required for the recovery from inactivation (Fig. 5).

By demonstrating that the wedge docks to a site that is enriched in binding sites for numerous PIEZO1 modulators (Fig. 2B) and considering that the wedge is encoded by a single exon (exon 32), the equivalent of which from PIEZO2 undergoes cell-type-specific splicing ^62^, our work also provides a framework for investigating the source of cell-type specific differences in PIEZO1 function that adjust its responsiveness to ensure optimal operation in different cellular environments.

## RESOURCE AVAILABILITY

### Lead Contact

Information and requests for resources and reagents should be directed to and will be fulfilled by the lead contact, Stefan G. Lechner.

### Material availability

The PIEZO1 plasmids generated in this study are available from the lead contact upon request.

### Data and code availability

Original Matlab code used in this paper to analyse Minflux images is available at GitHub (https://github.com/StefanLechnerUKE/PIEZO1_wedge_analysis). Raw data to generate and re-analyze Minflux experiments is also available in the corresponding GitHub repository. All data supporting the article, such as the MINFLUX and patch-clamp analysis outputs for each experimental condition and figures, are provided as source data. Any additional information required to re-analyse the data reported in this paper is available from the lead contact upon request.

## ACKNOWLEDGEMENTS

We thank Dr. Antonio Failla and the team of the imaging core facility at UKE Hamburg (DFG Research Infrastructure Portal #RI_00489) for technical assistance with the MINFLUX microscope. Funding for the MINFLUX microscope was awared by the the Hamburgische Investitions-und Förderbank (IFB, grant no. 51164232) under the Operational Programme Hamburg ERDF 2014-2020, REACT-EU of the European Regional Development Fund (ERDF). We also thank Ms. Kirsten Pfeiffer-Drenkhahn for assistance with calcium imaging experiment and Mr. Haider Al-Marsoomi as well as Ms. Claudia Lüchau for assistance with cloning and cell culture. This work was funded by the DFG grant LE3210/3-3 awarded to SGL.

## AUTHOR CONTRIBUTIONS

L.R. performed imaging and patch clamp experiments, wrote code and analyzed the data. C.V. performed Minflux and patch-clamp experiments and wrote analysis code. N.Z. performed patch-clamp experiments. S.G.L. conceptualized the study, acquired funding, wrote analysis codes, analyzed data and wrote the manuscript.

## SUPPLEMENTAL INFORMATION

Document S1, Figures S1-S4

## METHODS

### Experimental model

Mouse neuroblastoma Neuro-2a PIEZO1-Knockout cell line (N2A-P1KO) was generated previously from Neuro-2a ATCC CCL-131 (a gift from G.R. Lewin XX). Cell culture conditions were 37 °C atmosphere with 5% CO2 in a 1:1 mixture of Dulbecco’s Modified Eagle Medium and Optimal Minimal Essential Medium, with 10% Fetal Bovine Serum, 2 mM L-glutamine and 1% peniciline/streptomycine (all purchased from ThermoFisher).Seeding was performed on methanol– and acid-washed glass coverslips and coated with poly-Llysine (PLL, Sigma). 12 mm diameter coverslips were used for patchclamp recordings and calcium imaging, and #1.5 and 18 mm diameter for Minflux imaging. Tranfection was performed 24 to 48 hours after seeding with polyethylenimine (PEI, Linear PEI 25 K, Polysciences). For each mL of culture medium 7 µL with 29 µL of PBS and 0.6 µg of Plasmid DNA and incubated for 10 to 15 minutes at room temperature. The solution is then added to the culture medium and mixed gently by swirling. When preparing for calcium imaging experiments, 0.3 µg/mL medium of jGCaMP8m is added as well. After incubation for 24 hours, the medium is exchanged for fresh one and the cells are used after at least 24 and no later than 48 hours.

### Method details

#### Interdomain interaction predition

Prediction of interdomain interactions was performed by the RING4.0 algorithm^43^. The wedge as well as surrounding structures of the flat PIEZO1 (PDB: 7WLU) was used. The predicted interactions were then sorted by quality of interaction and bonds stronger than Van der Waals interactions were chosen.

#### Structure modelling and visualization

A full length PIEZO1 was generated of the flat (PDB: 7WLU) and the curved (PDBs: 6BPZ, 6B3R, 5Z10) structures which were aligned and superimposed for visualization purposes. All modification operations and visualization were performed in PyMol (Schrodinger). Plasmid design and visualization was performed with SnapGene (version 7, Dotmatics). All the other data, graphics and schematics were elaborated and visualized in Matlab, IgorPro, Illustrator (Adobe) and GraphPad Prism (version 10, GraphPad Software).

#### Generation of PIEZO mutants

PIEZO1-ALFA (instertion if ALFA-tag after H86) plasmids, which are expressing mutated mouse PIEZO1 and were previously shown not to affect PIEZO1 function^19^, were used as the template to generate firstly the PIEZO1-ALFA-Halo (insertion of Halo-tag at the C-terminus). From this template all other mutants used in this work were generated. Point mutants were introduced by site directed mutagenesis (primers were purchased from Sigma, sequence, template and obtained constructs can be found in the key resource table). The PCR reactions were digested with Dpln (New England Biolabs, 37 °C, 1 h) and column purified with standard kits (PureLink from Invitrogen or NucleoSpin from Macherey-Nagel) before transformation in electrocompetent Dh5a bacteria (Invitrogen) and grown at 37 °C overnight. Deletions were generated by Gibson cloning.

#### Electrophysiology

PIEZO1 mutants were characterized using twofold patch-clamp assays either stimulating the cell via indentation (poking, whole-cell) or applying negative pressure pulses to the patch (stretch, cell-attach). PIEZO1-ALFA-Halo served as the control in all experiments. All recordings were acquired using room temperature buffers that were adapted for each use (see below). An EPC10 amplifier with Patchmaster software (HEKA Elektronik) was used for all recordings. The pipettes were pulled with a P-97 Flaming-Brown puller (Sutter Instrument) from borosilicate pipettes (Sutter instruments) (2–6 MΩ for whole-cell, 1.5–4.5 MΩ after fire-polishing for cell-attach). Cells were held at a holding potential of –60 mV in both assays. The responses to mechanical stimulation were recorded with a sampling frequency of 50 kHz and a filter with a 2.9 kHz low pass filter. For whole-cell patch-clamp recordings the intracellular buffer contained: 125 mM K-gluconate, 7 mM KCl, 1 mM MgCl2, 1 mM CaCl2, 4 mM EGTA, 10 mM HEPES (pH 7.3 with KOH). The bath solution contained: 140 mM NaCl, 4 mM KCl, 1 mM MgCl2, 2 mM CaCl2, 4 mM glucose and 10 mM HEPES (pH 7.4 with NaOH). Mechanical stimulation was applied by poking the cells with a firepolished glass pipette (tip diameter 2–3 μm) that was positioned opposite to the recording pipette, at an angle of approximately 45° to the surface. The cells were stimulated in series of 15 stimuli with increments of 0.8 µm. The poking pipette was positioned and moved during the stimulation using a piezo driven micromanipulator (Preloaded Piezo actuator P-840.20, Physik Instrumente) with a speed of 1 μm/ms. For cell-attached pressure-clamp recording the intracellular buffer contained: 130 mM NaCl, 5 mM KCl, 1 mM MgCl2, 1 mM CaCl2, 10 mM HEPES, 10 mM TEA-Cl (pH7.3 with NaOH). The bath solution contained: 140 mM KCl, 1 mM MgCl2, 2 CaCl2, 10 Glucose, 10 HEPES (pH7.4 with KOH). The patch was stimulated by lowering the intrapipette pressure for 500 ms per stimulus in series of 16 increments of –5 mmHg using the High-Speed Pressure Clamp device (HSPC; ALA Scientific Instruments). A prepulse +5 mmHg was applied for 1 s before the stimulus to improve recovery from inactivation^37^. For repeated stimulations the cells were stimulated in series of 6 pulses, each lasting 250ms with –60 mmHg and +5 mmHg applied in the inter-pulse intervals. The duration of the inter-pulse intervals was varied between series applied consecutively to the same patch starting at 1000 ms and reducing the duration of by half until 50 ms were applied in the last series of six pulses. For prepulse-conditioned measurements the baseline amplitude was measured with a pulse of 500 ms at –60 mmHg following 500 ms at +5 mmHg, then holding the cell at 0 mmHg for 5 s to ensure full recovery and then giving a prepulse of –20 mV for 500ms before again stimulating with –60 mmHg. For the acquisition of I/V and single-channel conductance experiments, the stimulation pressure was adjusted for the individual cell to optimize for the evocation of single-channel openings. The conductance was determined by applying holding potentials from –140 mV to –40 mV in increments of 20 mV.

All electrophysiology was analysed using custom scripts in IgorPro (Wavemetrics). For poking-evoked currents, mechanical activation thresholds were defined as the stimulus that evoked the first peak current that was more than six standard deviations of the baseline above the baseline. The inactivation time constants (τinact) were measured by fitting the mechanically activated currents with a single exponential function (C1 + C2*e (–(t–t0)/τinact), where C1 and C2 are constants, t is time and τinact is the inactivation time constant (only peak currents between 100 and 1500 pA were used for these calculations and averaged per cell.

The peaks of pressure-induced currents (I) were normalized to the absolute maximal response of the cell at any pressure (Imax). Normalized pressure-response curve (I/Imax) from individual cells were fitted with a Boltzmann sigmoid to determine individual P50 (in mmHg). For repeated stimulations the peak of the ratio of the first peak and every subsequent peak (I/I_stim#1_) was calculated. When conditioning the cell with a prepulse, the ratio of the conditioned and unconditioned peak of the response to the –60 mmHg stimulation was calculated (I_+pre_/I_-pre_).

Single-channel amplitudes at a given holding potential (−140 mV to −40 mV, 20 mV steps) were determined as the difference between the peaks of the Gaussian fits of the trace histogram over multiple 1 s segments. Unitary conductance was determined from the linear regression fits of the I/V plot of individual cells. Recordings with excessive leak currents or unstable baseline were excluded. Recordings that displayed non-inactivating responses or unstable openings were also not used for further I/V analyses.

#### Calcium imaging

N2A-P1KO cells were cultured and transfected as described above. 48 to 72 hours after transfection the cells were washed once with PBS and incubated with a calcium imaging buffer containing 140 mM NaCl, 4 mM KCl, 1 mM MgCl2, 3 mM CaCl2, 4 mM glucose and 10 mM HEPES (pH 7.4 with NaOH). Fluorescent images were acquired every two seconds (500 ms exposure time) on an Olympus BX40 upright microscope equipped with standard Quad filter (Chroma), fluorescent lamp (HBO 100) and shutter (Lambda 10-2, Sutter Instrument) with a 20× water-immersion objective (UMPLFLN 20XW, Olympus), visualized with a Kiralux CS895MU camera (Thorlabs) and acquired with MicroManager ^63^. For perfusion and fast solution exchange a gravity-driven perfusion system (ValveLink8.2, AutoMate Scientific). The cell perfusion protocol was as follows: 30s wash with buffer, 30 s perfusion with Yoda2 solution (Tocris, 1 nM, 10 nM, 100 nM, 1 µM, 10 µM), 240 s wash with buffer. Two fields of view were acquired per coverslip moving for the second field of view upstream from the first, always using low Yoda2 concentrations (1 nM, 10 nM) as first and high Yoda2 concentrations (100 nM, 1 µM, 10 µM) as second stimulation to avoid desensitization.

To segment the cells and measure the fluorescence intensities a custom macro in ImageJ FIJI was used. The ratio of the fluorescence and the baseline (mean fluorescence of the first 15 s of the initial wash with buffer) was calculated over time (F/F_0_). Within the Yoda2 perfusion time, the maximal F/F_0_ per cell was extracted and fitted with a sigmoid curve for EC_50_ calculation.

#### Preparation of samples for 3D-MINFLUX/DNA-PAINT imaging

N2A-P1KO cells were plated on 18mm coverslips and transfected as described above; the cells were prepared for imaging 48 to 72 hours after transfection. Preparation started by fixing the cells in paraformaldehyde (1%) and glutaraldehyde (0.05%) and quenched for 10 min 5 mM NaBH4 and then with a mix of 50 mM glycine and 50 mM NH4Cl, all diluted in PBS and washing with PBS. Samples were blocked with Antibody Incubation buffer (Massive Photonics) for 30 min at room temperature. The cells were incubated for 1 h at room temperature or overnight at 4 °C with Massive-Tag-Q Anti-ALFA nanobodies conjugated with a DNA docking strand (both from Massive Photonics) diluted at 1:100 (Anti-ALFA) in Antibody Incubation buffer. After washing thrice with 1× Washing buffer (Massive Photonics), the cells were post-fixed for 5 min and the fixatives were quenched and washed as described above. Samples were then incubated for 10 min with 200 μl of gold nanoparticles for future stabilization (gold colloid 250 nm, BBI Solutions). The cells were washed several times with PBS to remove excess nanoparticles and remaining nanoparticles were stabilized by incubating the sample with PLL for 60 min at room temperature. Cells were again washed thrice with PBS before mounting and used over the course of the following two days.

Corresponding DNA-PAINT imagers (Massive Photonics) conjugated to Atto 655 were freshly diluted in Imaging buffer (Massive Photonics) for a final concentration of 1 ∼ 2 nM (Atto 655, Imager sequence #3). A drop of imager dilution was added into a cavity slide and coverslips were mounted and sealed with Picodent Twinsil (Picodent).

#### 3D-Minflux/DNA-PAINT imaging

An Abberior MINFLUX commercial microscope built on an inverted IX83 microscope with a 100× UPlanXApo objective (Olympus) and using Imspector Software (Abberior Instruments) was used to acquire all Minflux data. The calibration and alignment of the excitation beam pattern as well as the pinhole positioning were performed at the beginning of each acquisition session using fluorescent nanoparticles (Abberior Nanoparticles, Gold 150 nm, 2 C Fluor 120 nm, Abberior Instruments). Cells transfected with PIEZO1-Mutants were identified with a 640 nm confocal scan. Focus was set to the interface between coverslip and cell membrane. At least 2 gold fiducials were present in the field of view and used by the activefeedback stabilization system of the microscope (IR 975 nm laser, Cobolt, and CCD camera, The Imaging Source), resulting in a precision below 1 nm in all three axes and being stabled for hours. A ROI of approximately 3 × 3 to 5 × 5 μm (and up to 8 × 8 μm for some overnight recordings) was selected. Laser power in the first iteration was set at 16% laser power and pinhole was set to 0.83 A.U. Final laser power in the last iteration is scaled up by a factor of six. ROIs were imaged for at least 2 h and up to overnight (∼ 12 h) using the manufacturers default 3D Minflux TCP sequence. Detection for Atto-655 signals was performed with two avalanche photodiodes channels (650–685 nm and 685–720 nm) that were pooled.

#### 3D-Minflux data analysis

Valid Minflux localization data from the final iterations was exported as MATLAB files from the Imspector software and analyzed using costum matlab scripts for post-processing and filtering, further operations and data visualization was also performed in Matlab. The data was filtered in a stepped process firstly by cfr (center frequency ratio, directly implemented during the measurement, set at 0.8) and efo (effective frequency at offset, kHz, retrieved for each individual valid locations) to dismiss potential multiple emitters. Secondly localizations from the same emission trace (ones sharing one trace identification number (TID)), traces with a standard deviation higher than 10 nm in any dimension and such that contained less than 3 localizations were filtered-out. The first two localizations of each trace were discarded, since they tend to deviate from the majority of the cloud. A scaling factor of 0.7 was applied to all z-coordinates to correct for the refractory index mismatch between coverslip and sample ^48^. The geometric center for each individual filtered trace was then calculated. Due to the nature of DNA-PAINT, different traces originating from repeated detection of the same PIEZO protomer were identified using DBSCAN clustering with minPoints of 2 and epsilon of 8 nm. This cutoff was selected based on the localization precision of MINFLUX and the possible ALFA-tag flexibility ^18^. Protomers that were detected several times were represented by the mean coordinate of the localizations clustered by this DBSCAN algorithm. Lastly PIEZO1 trimers that were a) fully labelled and b) fully detected by Minflux were determined by searching the cloud of traces for clusters of three adjacent traces with distances to each other that were smaller than 40 nm (the assumed maximal distance two Atto-655 molecules bound to the same PIEZO1 trimer can possibly have based on available flattened PIEZO1 ^18^ and distances to the next closest trace of over 60 nm. Only trimers with a maximum internal angle of less than 120° were included. The mean interblade distance for each trimer was calculated. Further visualization of trimers was done by a 2D in-plane projection of the raw localizations for each protomer, followed by a fit with a bivariate Gaussian distribution and displayed with its probability density.

### Quantification and statistical analysis

#### Statistical tests and reproducibility

All experiments in this study were performed independently at least three times, yielding similar results. For Minflux experiments, data are from at least 4 cells from at least 3 independent experiments. No statistical method was used to predetermine sample size. Experiments were not randomized and investigators were not blinded during experiments and analysis. Data distribution was systematically evaluated using D’Agostino–Pearson test and parametric or nonparametric tests were chosen accordingly. The statistical tests that were used, the exact P-values and information about the number of replicates are provided in the figure or the corresponding legends.

