## Supplementary material for "State-dependent binding of the wedge domain controls inactivation of the mechanosensitive ion channel PIEZO1": Suppl Data

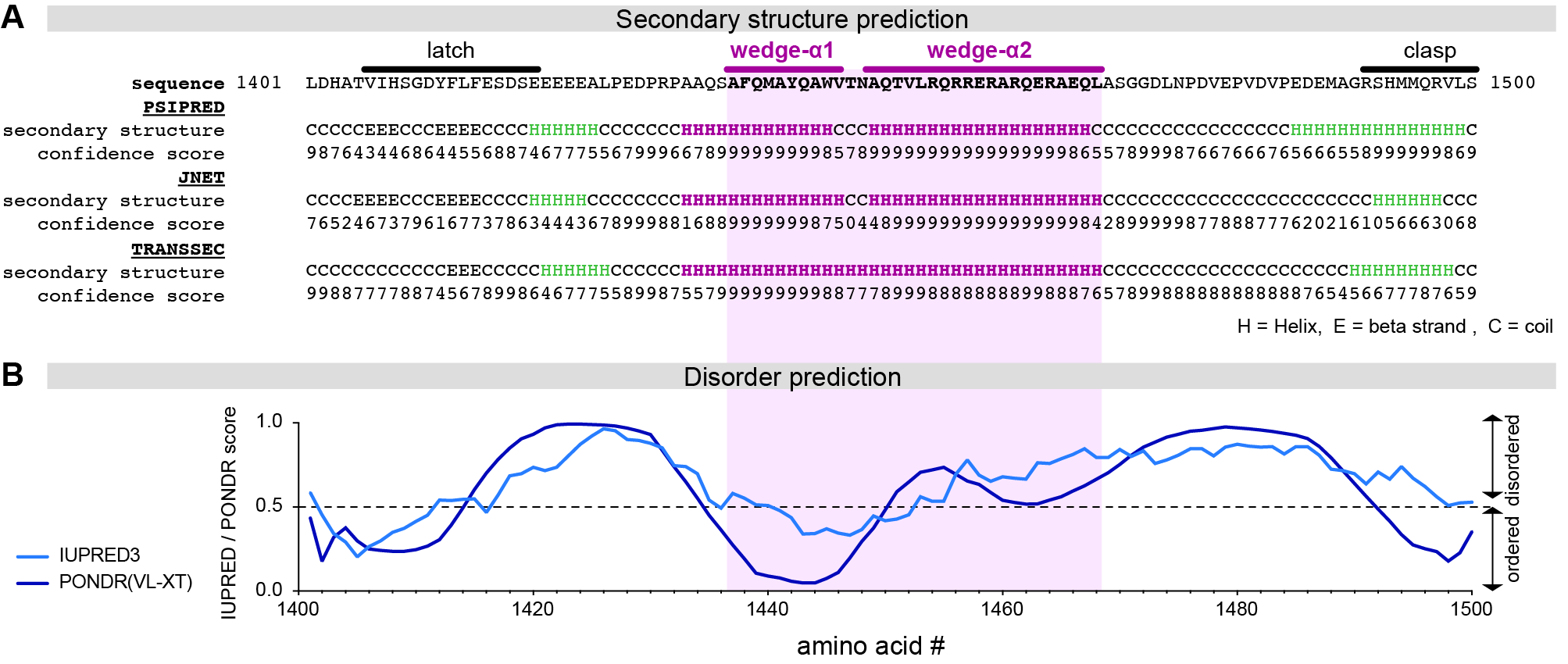

**Figure S1 | In-silico prediction of the secondary structure of the wedge and adjacent linkers**

**(A)** The indicated amino acid sequence (top) including the wedge and the adjacent regions was submitted to the PROTEUS structure predictino server 2.0, which implements PSIPRED, JNET and the locally developed TRANSSEC algorithm for secondary structure prediction. Predicted secondary structures are indicated by H(helix), E (beta strand) and C (coil) and PROTEUS confidence scores (1-10, with 10 indicating highest confidence). Note, all three algorithms predict a helical secondary strucutre for the wedge domain.

**(B)** The same amino acid sequence shown in (A) was analyzed for the presence of intrinsically disordered regions using the IUPRED3 (bright blue) and PONDR(VL-XT; dark blue) algorithms. Note, the wedge, specifically the a1-helix, has a very low disorder score, whereas the adjacent linkers appear to be highly diordered.

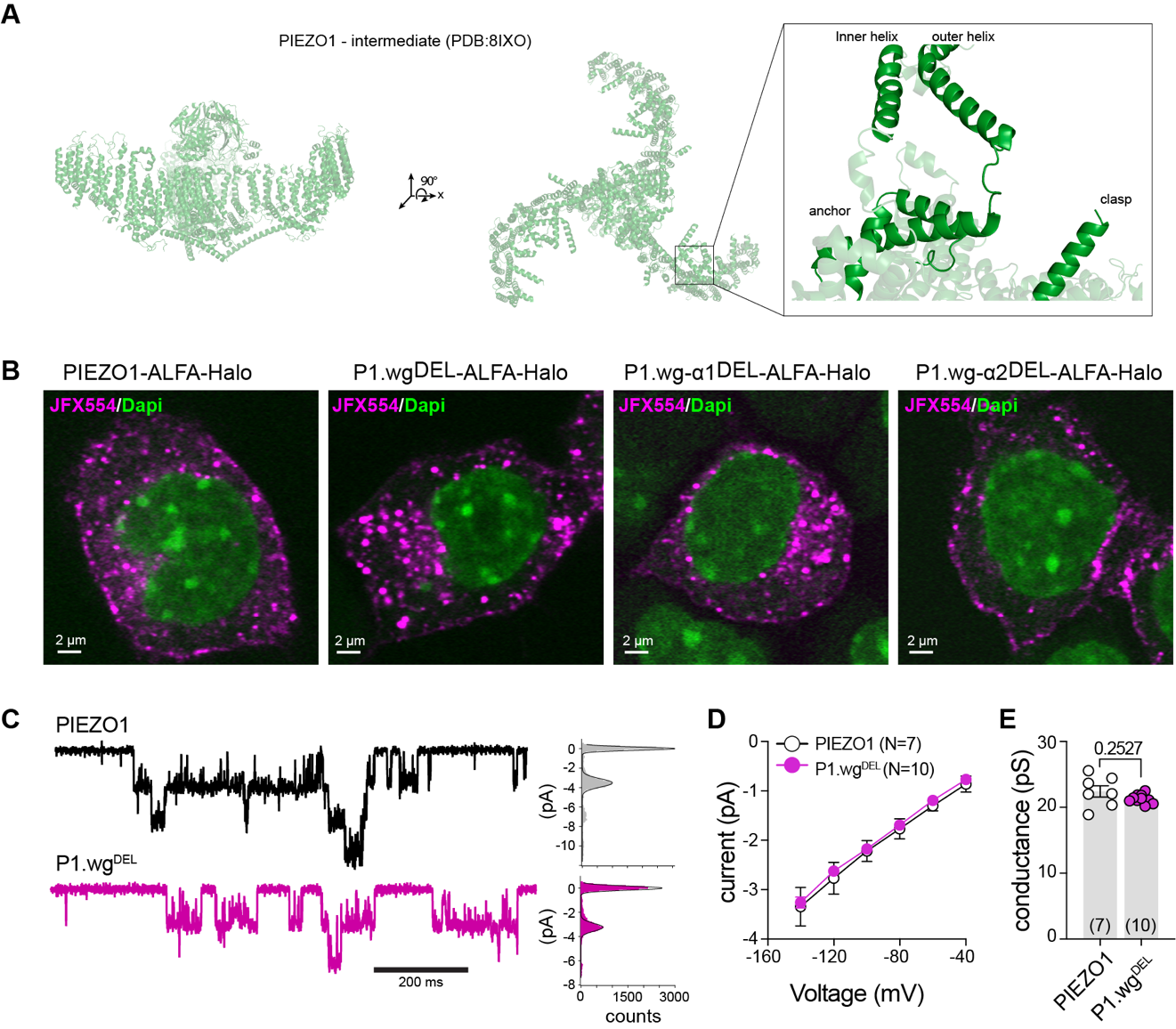

**Figure S2 | Characterization of PIEZO1 wedge deletion mutants**

**(A)** Side (left) and top (middle) view of the intermediate open structure of the PIEZO1-S2472E mutant. Close-up view (right) shows the absence of the wedge docking to anchor-OH linker.

**(B)** confocal images of the indicated PIEZO1 variants showing prominent plasma membrane expression.

**(C)** Example traces of stretch-evoked single channel openings of PIEZO1 (top left) and Wedge^DEL^ (bottom left), corresponding all-points histogram (right), with Gaussian fits and indicated peak average of PIEZO1and Wedge^DEL^.

**(D)** Current/voltage relationship of single-channel amplitude of PIEZO1 and P1.wgDEL fitted with a linear regression.

**(E)** Comparison of the mean ± s.e.m. single channel conductance, deduced from the data shown in (D), using Student’s unpaired t-test with Welch’s correction. N numbers are indicated inside the bars and values from individual cells are shown a cricles.

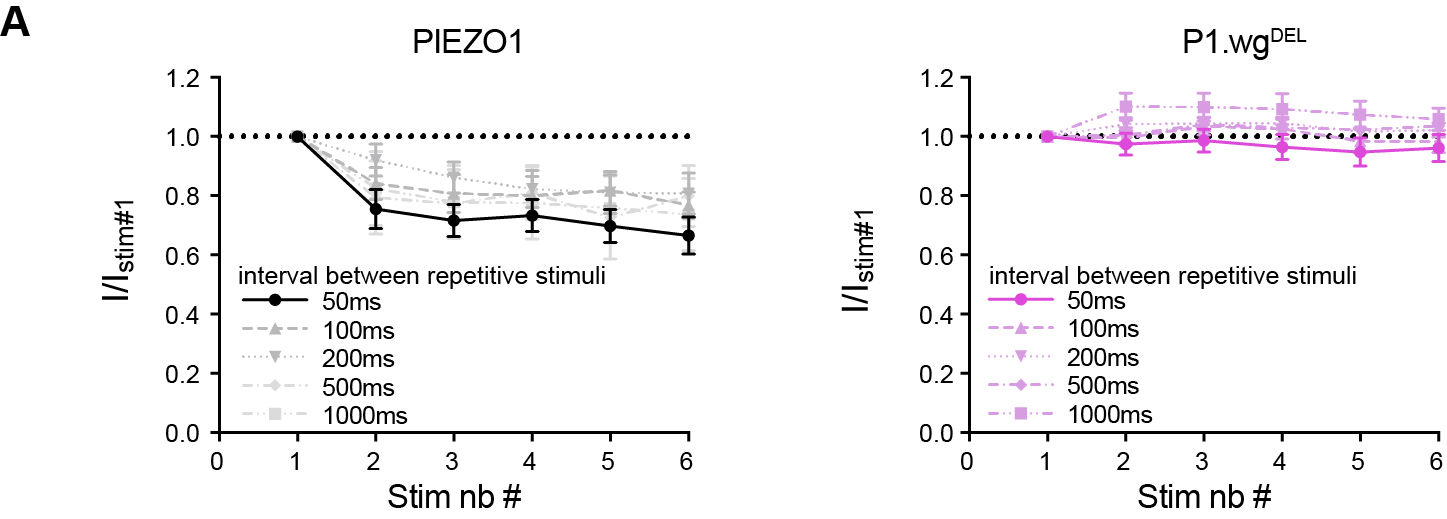

**Figure S3 | Recovery from inactivation of PIEZO1 and P1.wg^DEL^**

**(A)** Example traces of PIEZO1 (top left) and P1.wg^DEL^-mediated (bottom, left) currents elicited by repetitive mechanical stimulation. Magnitude of responses evoked by repeated mechanical stimulation (200 ms, -60 mmHg) at the indcated time intervals and normalized to the amplitude of the response evoked by the first stimulus in cells expressing PIEZO1 (left) and P1.wg^DEL^ (right). Symbols represent means ± s.e.m.

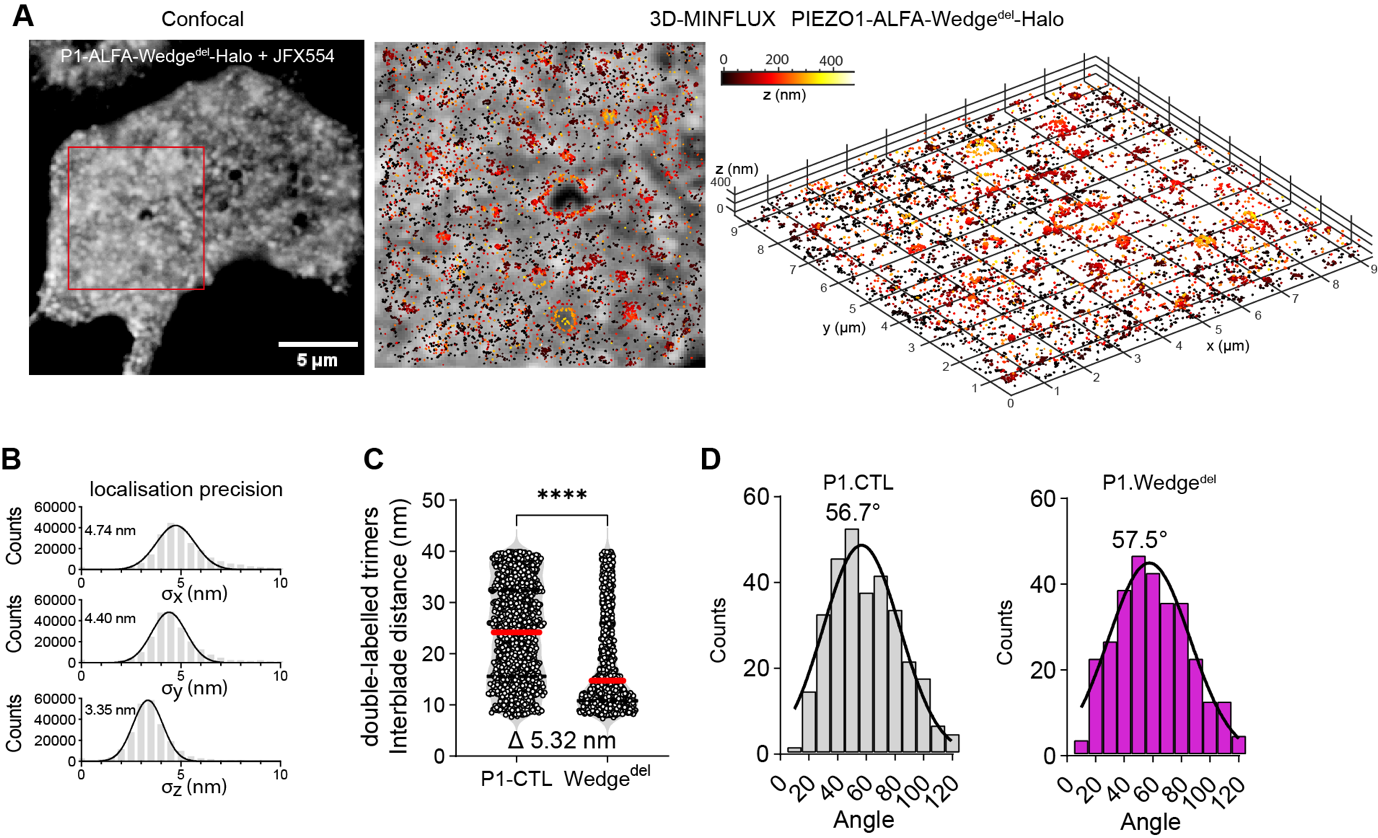

**Figure S4 | 3D-MINFLUX characterization of PIEZO1 and P1-wgDEL**

**(A)** Confocal image of P1.wg^DEL^ expressing N2a.P1KO cell (left), close-up view of region indicating by red square with superimposed MINFLUX raw data (middle), and perspective view of the MINFLUX data only from the same region.

**(B)** Histograms of standard deviations of trace locations in x (top), y (middle) and z (bottom) direction in nanometres and Gaussian fits. Respective µ-values of the Gaussian fits are indicated next to the peaks.

**(C)** Comparision of interblade distances measured in double-labelled PIEZO1 and P1.wg^DEL^ trimers using Mann-Whitney test (****, P< 0.0001). Medians are indicated as red lines, quartiles as dotted black lines and delta of means is indicated at the bottom.

**(D)** Histograms of the interblade angles (see Fig. 4B right panel for explanation) of the triple labelled PIEZO1 (left) and P1.wg^DEL^ (right) trimers with Gaussian fit and means indicated above peak.

### **Table S1**

| **Plasmid Name** | **Primer** | **Sequence** |
| --- | --- | --- |
| P1_ALFA_Halo | Vector_FWD | TAAAGGCGCTCCTGCCCTGGCAACCA |
|  | Vector_REV | GAGGTTCCATGGTGGCGCGGCC |
|  | Fragment_FWD | CCGCGCCACCATGGAACCTCACGTGCTCG |
|  | Fragment_REV | CCAGGGCAGGAGCGCCTTTAAGTCCCACAG |
| P1_ALFA_Halo_Y1442A | FWD | CCTTCCAGATGGCAGCCCAGGCATGGGTAACCAATGC |
|  | REV | GCATTGGTTACCCATGCCTGGGCTGCCATCTGGAAGG |
| P1_ALFA_Halo_W1445A | FWD | CCAGATGGCATACCAGGCAGCGGTAACCAATGC |
|  | REV | GCATTGGTTACCGCTGCCTGGTATGCCATCTGG |
| P1_ALFA_Halo_R1456A | FWD | GACAGTGCTGAGGCAGGCGCGGGAGCGGGCACGG |
|  | REV | CCGTGCCCGCTCCCGCGCCTGCCTCAGCACTGTC |
| P1_ALFA_Halo_Y2175A | FWD | GACAGAGAAGAAAGCCCCCCAGCCCAAGG |
|  | REV | CCTTGGGCTGGGGGGCTTTCTTCTCTGTC |
| P1_ALFA_Halo_K2184A | FWD | GGGGCAGAAGAAGGCGAAAATTGTCAAGTATGG |
|  | REV | CCATACTTGACAATTTTCGCCTTCTTCTGCCCC |
| P1_ALFA_Halo_Y1442A-W1445A | FWD | GAGTGCCTTCCAGATGGCAGCCCAGGCAGCGGTAACC |
|  | REV | GGTTACCGCTGCCTGGGCTGCCATCTGGAAGGCACTC |
| P1_ALFA_Halo_wg^DEL^ | Vector_FWD | CCTGCATATCCCTGAGCTGGAG |
|  | Vector_REV | TCAAGTCACCCTGAGCTGCAGGCCTGG |
|  | Fragment_FWD | TGCAGCTCAGGGTGACTTGAACCCAGATGTGGAAC |
|  | Fragment_REV | CCAGCTCAGGGATATGCAGGCG |
| P1_ALFA_Halo_wgα2^DEL^ | Vector_FWD | CCTGCATATCCCTGAGCTGGAG |
|  | Vector_REV | TCAAGTCACCATTGGTTACCCATGCCTGGTATG |
|  | Fragment_FWD | GGTAACCAATGGTGACTTGAACCCAGATGTGGAAC |
|  | Fragment_REV | CCAGCTCAGGGATATGCAGGCG |
| P1_ALFA_Halo_wgα1^DEL^ | Vector_FWD | CCTGCATATCCCTGAGCTGGAG |
|  | Vector_REV | CTGTCTGGGCCTGAGCTGCAGGCCTGG |
|  | Fragment_FWD | TGCAGCTCAGGCCCAGACAGTGCTGAGGCA |
|  | Fragment_REV | CCAGCTCAGGGATATGCAGGCG |
| P1_ALFA_Halo_Y2175A_K2184A | FWD | GACAGAGAAGAAAGCCCCCCAGCCCAAGG |
|  | REV | CCTTGGGCTGGGGGGCTTTCTTCTCTGTC |
